# GAT-FD: An Integrated MATLAB Toolbox for Graph Theoretical Analysis of Task-Related Functional Dynamics

**DOI:** 10.1101/2021.02.21.432161

**Authors:** Meng Cao, Ziyan Wu, Xiaobo Li

**Affiliations:** Department of Biomedical Engineering, New Jersey Institute of Technology, NJ, USA; Department of Electrical and Computer Engineering, New Jersey Institute of Technology, NJ, USA

**Keywords:** Task-based fMRI, Graph Theoretical Techniques, Dynamic Functional Connectivity, Network Analysis, GAT-FD.

## Abstract

Functional connectivity (FC) has been demonstrated to be varying over time during sensory and cognitive processes. Quantitative examinations of such variations can significantly advance our understanding on large-scale functional organizations and their topological dynamics that support normal brain functional connectome and can be altered in individuals with brain disorders. However, toolboxes that integrate the complete functions for analyzing task-related brain FC, functional network topological properties, and their dynamics, are still lacking. The current study has developed a MATLAB toolbox, the Graph Theoretical Analysis of Task-Related Functional Dynamics (GAT-FD), which consists of four modules for sliding-window analyses, temporal mask generation, estimations of network properties and dynamics, and result display, respectively. All the involved functions have been tested and validated using fMRI data collected from human subjects when performing a block-designed task. The results demonstrated that the GAT-FD allows for effective and quantitative evaluations of the functional network properties and their dynamics during the task period. As an open-source and user-friendly package, the GAT-FD and its detailed user manual are freely available at https://www.nitrc.org/projects/gat_fd and https://centers.njit.edu/cnnl/gat_fd/.

## 1. Introduction

Functional connectivity (FC), which quantifies temporal dependencies among spatially separated brain regions, has been highlighted as a sensitive and robust measurement in functional magnetic resonance imaging (fMRI) for understanding the topological organization of functional brain networks during sensory and cognitive processes and at resting-state (Power et al., 2010;Du et al., 2018). Typically, FC was evaluated in a “static” sense. Recently, accumulative evidence has suggested the temporally varying pattern of FC, referred to as dynamic FC, which can provide us a novel approach to depicting the non-stationarity of functional brain communications (Horovitz et al., 2008;Bassett et al., 2011;Fornito et al., 2012).

Currently, sliding-window-based techniques are commonly implemented for estimating FC dynamics (Sakoglu et al., 2010;Hutchison et al., 2013;Rashid et al., 2014;Preti et al., 2017). Such approach applies an N-point moving window along the time domain to estimate the pair-wise FCs at the current time point based on signals of the current and previous N time points. It thus generates a series of consecutive FC metrics, depicting the dynamically varying FC and topological organizations of the functional brain network. To date, several toolkits have been proposed for sliding-window analysis, including GIFT (http://mialab.mrn.org/software/gift/) (Allen et al., 2014), DyNaConn (https://bitbucket.org/johnesquivel/dynaconn/src/master/), DyConPro (https://github.com/tobiamj/DyConPro) (Tobia et al., 2017), DynamicBC (https://www.nitrc.org/projects/dynamicbc/) (Liao et al., 2014), and CONN (https://web.conn-toolbox.org/) (Whitfield-Gabrieli and Nieto-Castanon, 2012). Although these packages have made it possible to generate sliding window-based connectivity matrices, they either were designed specifically for resting-state FC dynamics analysis or require extra tools for calculating topological dynamics of the functional brain network. Moreover, none of these existing toolboxes provide functions for quantification of the dynamics associated with the FC and network topology.

In this study, we introduce an open-source and user-friendly MATLAB toolbox, the Graph Theoretical Analysis of Task-Related Functional Dynamics (GAT-FD), which we have developed to integrate the complete pipelines for estimating the task-related dynamic brain FC and quantifying the topological dynamics and its statistical property of the functional brain network. This toolkit and the user manual with detailed explanations of each functions and step-by-step instructions for implementations are freely available at https://www.nitrc.org/projects/gat_fd and https://centers.njit.edu/cnnl/gat_fd/.

## 2. Methods

### 2.1. Overview

The GAT-FD toolbox provides a graphical user interface for characterizing the functional network dynamics in task-related fMRI data, based on the graph theoretical techniques. It was developed in MATLAB (Mathwork, Inc.) version 2019b and has been tested with MATLAB version 2018b to version 2019b. The GAT-FD is organized into four modules (**Figure 1A**) for sliding-window analysis (**Figure 1B**), task-specific temporal mask generation (**Figure 1C**), estimations of network properties (**Figure 1D**), and result display (**Figure 1E**), respectively.

**Figure 1.**
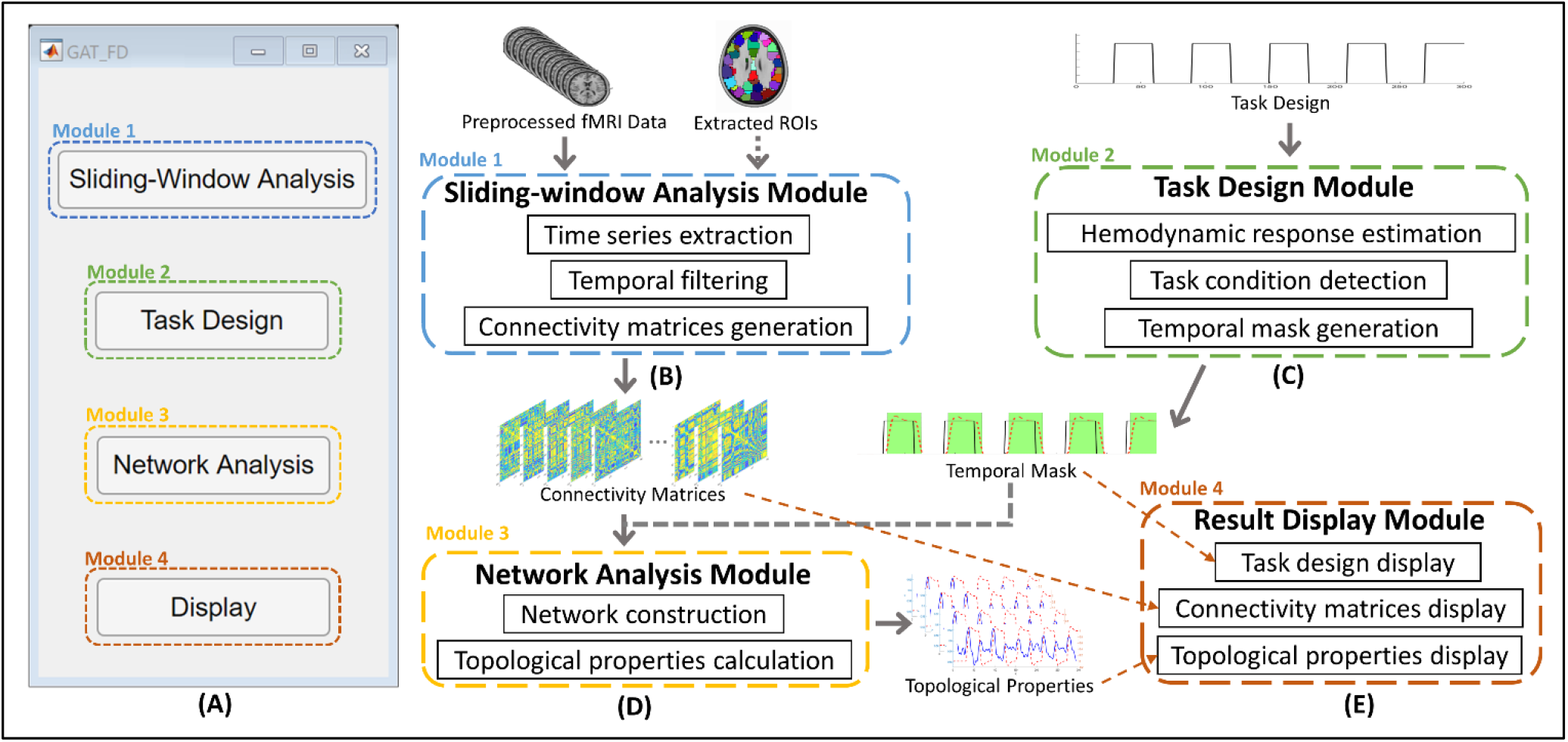
The four function modules of the GAT-FD toolbox. (A) The main user interface. (B) The sliding-window analysis module. This module generates connectivity matrices based on specified sliding-window parameters. (C) The task design module. This module generates temporal inclusive mask based on task design. (D) The network analysis module. This module calculates the topological properties for each connectivity matrix. (E) Result display module. This module provides visualization of the results from all other modules. fMRI: functional magnetic resonance imaging. ROIs: regions-of-interest. Note: Required steps are indicated using solid arrows. Optional steps are indicated using dashed arrows.

### 2.2. Inputs

The GAT-FD toolbox works with pre-processed fMRI data. Properly pre-processed images can effectively minimize the falsely discovered dynamics caused by motion artifacts and undesired physiological fluctuations. The inputs are 4-dimonsional fMRI data in uncompressed or compressed NIfTI format (e.g., *.nii or *.nii.gz). All build-in atlases are in Montreal Neurological Institute (MNI) space. Therefore, all input data are required to be transformed from individual imaging space to MNI space, if the user is intended to use any of the build-in atlas. The toolbox also provides an option for the user to import MAT-format input files with a matrix containing the time series of each region-of-interest (ROI), which allows extra flexibility in temporal processing.

### 2.3. Sliding-window analysis and FC matrix construction

The sliding-window approach is the most common analytical method to explore the network dynamics in fMRI studies (Preti et al., 2017;Gonzalez-Castillo and Bandettini, 2018). In this approach, a sliding-window with a fixed length (called as window size) and a “moving step” (called as step size) along the time series are first defined. A FC matrix is constructed based on the current time point and the previous ones which are covered by the sliding-window. Then the sliding-window moves to next time point and the process repeats till the end of the task period to generate a series of FC metrices. The GAT-FD toolbox includes a sliding-window analysis module to extract the activation time series for selected ROIs, perform temporal filtering, apply sliding-window, and construct the FC matrices, as shown in **Figure 2**. Two options for ROI determination, the customized brain masks and build-in atlas, are available in the module. The toolbox currently provides two build-in atlas, including the automated anatomical labeling (AAL) atlas (Tzourio-Mazoyer et al., 2002) and Brainnetome atlas (Fan et al., 2016). For user-loaded (customized) brain masks and atlas, the MNI space format is required to avoid miscalculations during extraction of blood-oxygen-level-dependent (BOLD) responses.

**Figure 2.**
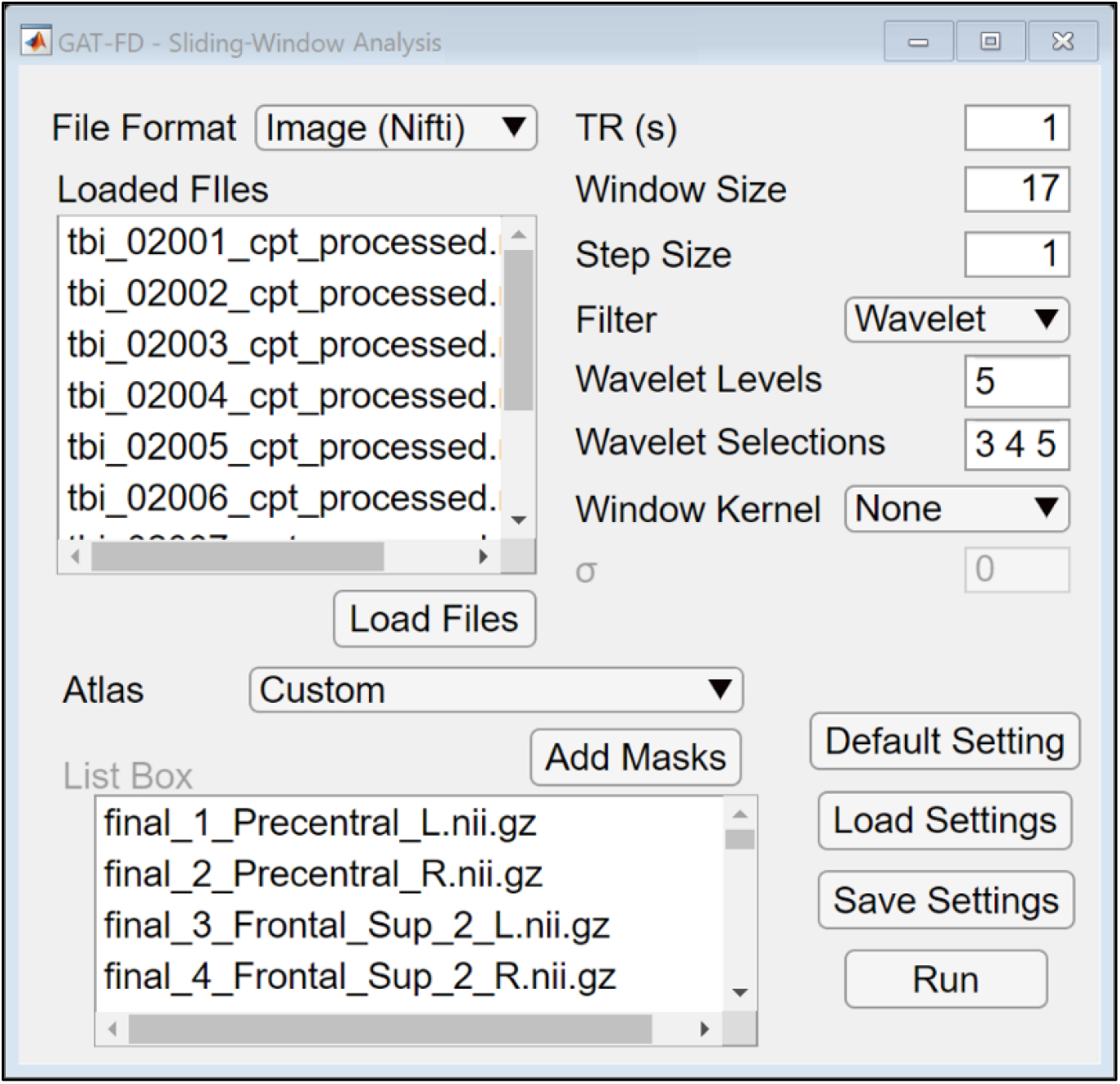
The user interface of sliding-window analysis module. By clicking the “Default Setting” button, users can reset all the parameters to default values. The user specified parameters can be saved and loaded by clicking the “Save/Load Settings” buttons. By clicking the “Run” button, the sliding-window analysis can be performed.

Compared to static FC analysis that uses signals in the entire task duration to calculate the functional correlations, dynamic FC analysis uses a much smaller window size, which becomes more sensitive to noises in the data. Therefore, the step of temporal filtering is essential to guarantee the accuracy of detected dynamics of the functional brain network. The sliding-window analysis module of the GAT-FD toolbox includes options of high-, low-, band-pass filter, and wavelet filter to help further minimize the undesired noises. When applying high-, low-, or band-pass filter, the corresponding cut-off values (a lower-limit cut-off value is required for high-pass filter, an upper-limit cut-off value is required for low-pass filter, and both lower- and upper- limits cut-off values are required for band-pass filter) are specified as the inverse of the cut-off frequency, i.e., the unit of second, instead of Hz are implemented here. Wavelet decomposition, as a frequently used tool in task-based fMRI data analysis, increases sensitivity in detecting signal correlation against a noisy background, especially when motion artifacts related spikes occur (Bullmore et al., 2004;Ginestet and Simmons, 2011). If the wavelet filtering function is selected, the sliding-window module will first decompose each activation time series using the maximal overlap discrete wavelet transform (MODWT) with a specific number of levels, and then transfer back with selected levels of coefficients using inverse MODWT. The number of wavelet decomposition levels and the selected wavelet levels need to be specified by the user. The temporal filtering function is optional in this module, given that input data can be already filtered during the pre-processing steps.

In the GAT-FD toolbox, the FC at the current time point between a pair of ROIs is represented by the Pearson’s correlation coefficient of the BOLD signals within the corresponding sliding-window in the two ROIs. Therefore, the window size and the step size are critical in detecting the desired temporal dynamics during the task when using sliding-window analytical method. The selections of these parameters depend on the task design and the repetition time (TR). Studies have suggested the window size to be larger than 15 TRs to get reliable estimations of between-region temporal correlations (Braun et al., 2015;Di et al., 2015;Rosenthal et al., 2017). In addition, the window size needs to be smaller than the length of one task block to provide sufficient number of measures on describing mid-task variability (Li et al., 2018;Betzel et al., 2020). The GAT-FD toolbox also offers an option to utilize gaussian kernel-based sliding-window (Allen et al., 2014;Preti et al., 2017). Such approach applies a tapered sliding-window with a different weight for each time point involved in the window. A series of FC matrices is then generated for each subject with user specified sliding-window parameters. The sliding-window analysis module is able to process multiple input files at once, to generate one MAT-format output file for each selected subject.

### 2.4. Temporal inclusive mask generation for block-designed tasks

Studies have found that the pattern of pairwise FC varies between the rest and task conditions during fMRI (Spadone et al., 2015;Shah et al., 2016). To map the task-specific (or rest-specific) FC matrices in the sliding-window analysis module, the temporal inclusive mask generated from the task design module of the GAT-FD toolbox can be needed. To determine the temporal inclusive mask, two thresholding methods, estimated activation level thresholding and condition coverage percentage thresholding, are provided (**Figure 3)**. Users can choose to use either or both methods to generate their study-specific masks by checking the box of each function. If both boxes are checked, logic AND will be implemented to the two masks for the final output of the temporal inclusive mask. The estimated activation level thresholding method is based on the estimated hemodynamic responses which are internally estimated in the module by convolving the task design with the hemodynamic response function in Statistical Parametric Mapping (SPM) toolbox (Penny et al., 2011). If the box for estimated activation level method is checked and the threshold is provided by the user (or by using default value), the time points with the estimated activation magnitude (ranged from 0 to 1, with 1 representing the maximum estimated response for a single stimulus) higher than the user defined threshold are included in the temporal inclusive mask. By checking the box of condition coverage percentage thresholding method, selecting the condition type (1 for task and 0 for rest), and defining the percentage threshold (X%), time points with at least X% of their associated sliding windows under the selected task condition (according to the task design) will be inclusive in the mask. The default settings for these two parameters are 0.8 and 80, respectively. The output of this module is a temporal inclusive mask which contains the indexes of the task-related study-specific FC matrices.

**Figure 3.**
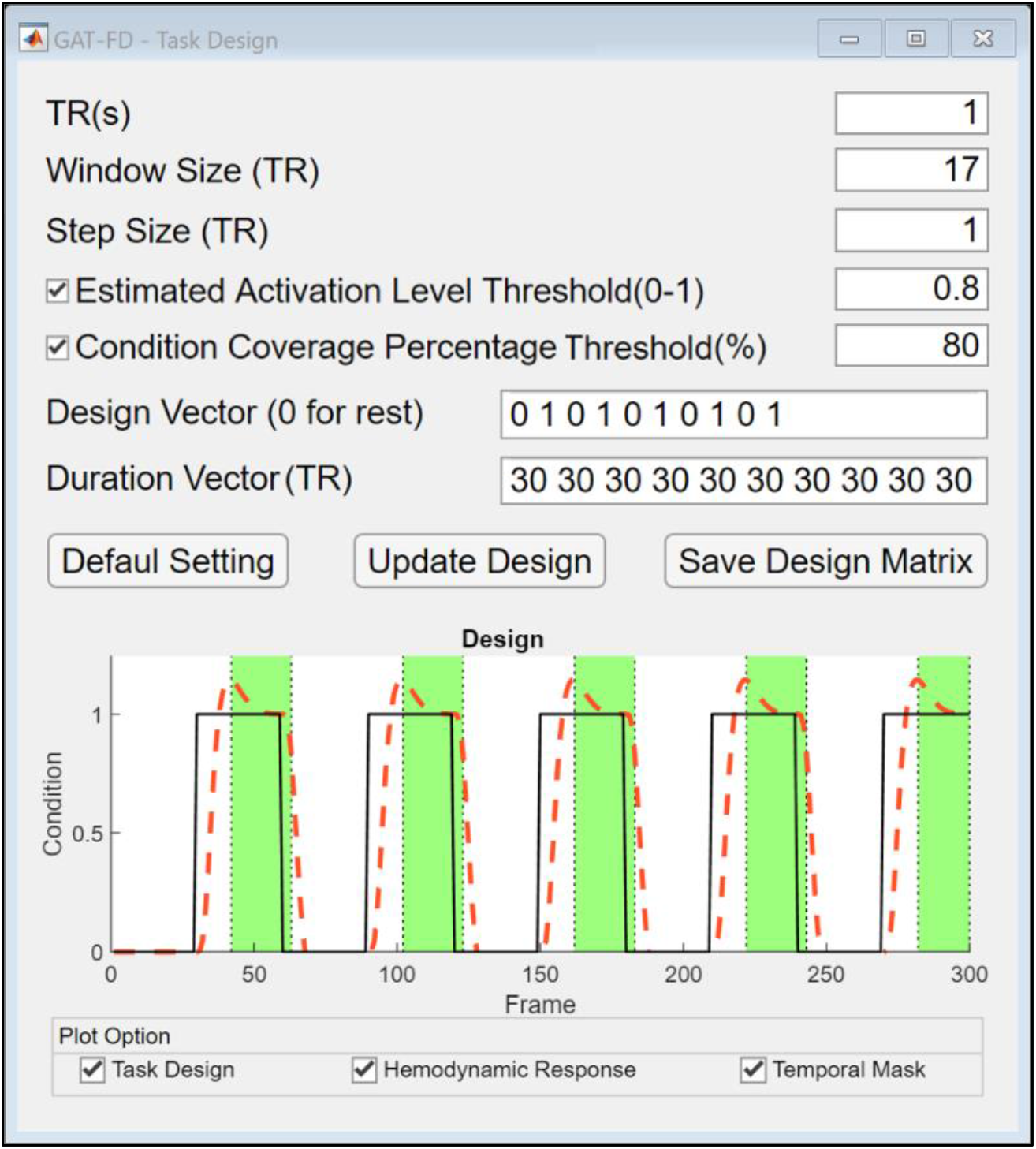
The user interface of task design module. By clicking the “Default Setting” button, users can reset all the parameters to default values. After defining all the parameters, the temporal inclusive mask can be plotted in the bottom by clicking the “Update Design” button. When the “Task Design” plot option is checked, the task design is plotted in solid black line. When the “Hemodynamic Response” plot option is checked, the estimated hemodynamic response is plotted in dashed red line. When the “Temporal Mask” plot option is checked, the time points selected by the temporal inclusive mask are shown in green areas. The temporal inclusive mask file needs to be saved by clicking on the “Save Design Matrix” button.

### 2.5. Characteristics of the topological dynamics of the functional brain network

To characterize the topological dynamics of the functional brain network, the FC matrices need to be binarized. The network analysis module of the GAT-FD toolbox provides multiple thresholding methods for FC matrix binarization, including absolute thresholding, proportional thresholding, and wiring cost thresholding. The absolute thresholding method applies the same user specified correlation coefficient threshold (ranging from −1 to 1) to all connectivity matrices. The proportional thresholding method defines the threshold relative to the maximum correlation coefficient in a FC matrix with user specified proportion, ranging from 0 to 1. The wiring cost thresholding method preserves the top connections (with user specified cost threshold) in the connectivity matrix. The cost of a network is defined as the number of existing connections divided by the number of all possible connections, which ranged from 0 to 1, representing the top 0% to 100% connections, respectively.

Next, the network topological properties at both global and nodal levels are calculated based on the constructed binary-networks within and averaged over the specified threshold range. The global metrics include network global efficiency, network local efficiency, network averaged degree, and network averaged clustering coefficient. The nodal metrics include nodal global efficiency, nodal local efficiency, nodal degree, nodal clustering coefficient, and betweenness centrality. The variance of each topological property over the selected timepoints can be calculated when the temporal inclusive mask file from task design module is provided. The variance is defined as, 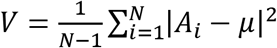, where N is the number of timepoints, *A_i_*; is the network topological property at that time points, and *μ* is the mean of the network topological property over the selected timepoints. The GAT-FD toolbox utilizes functions from the brain connectivity toolbox for network construction and topological features calculation (Rubinov and Sporns, 2010). The calculations are time consuming, therefore, parallel computing option in this module is supported if the MATLAB parallel toolbox is installed. The detailed configurable parameters are shown in **Figure 4**.

**Figure 4.**
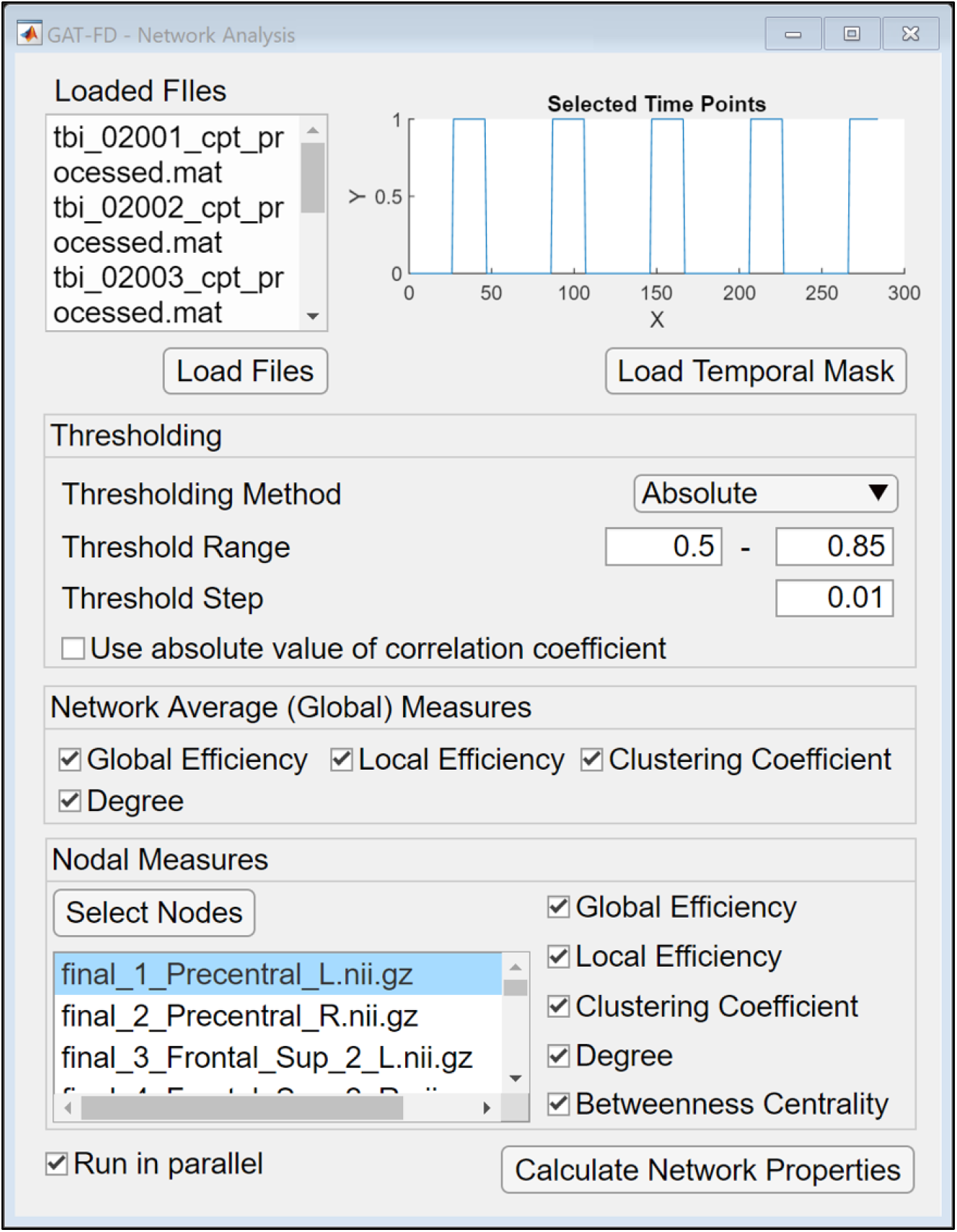
The user interface of network analysis module. The selected timepoints are displayed at the top right corner after loading the temporal mask file. If the “Use absolute value of correlation coefficient” is checked, all the negative correlation coefficients in the connectivity matrices are converted to positive value before thresholding. By clicking the “Calculate Network Properties” button, the network analysis is performed with user specified thresholding parameters and user selected topological measures.

### 2.6. Result Display

Result display module provides convenient features to visually check the calculated results from all previous modules, as shown in **Figure 5**. The constructed connectivity matrices for each sliding-window can be checked for any abnormal conditions. Task design can be loaded and displayed to provide a visual inspection for sliding-window selection. In addition, the calculated network topological properties can be displayed individually or as a group, along with the estimated hemodynamic response.

**Figure 5.**
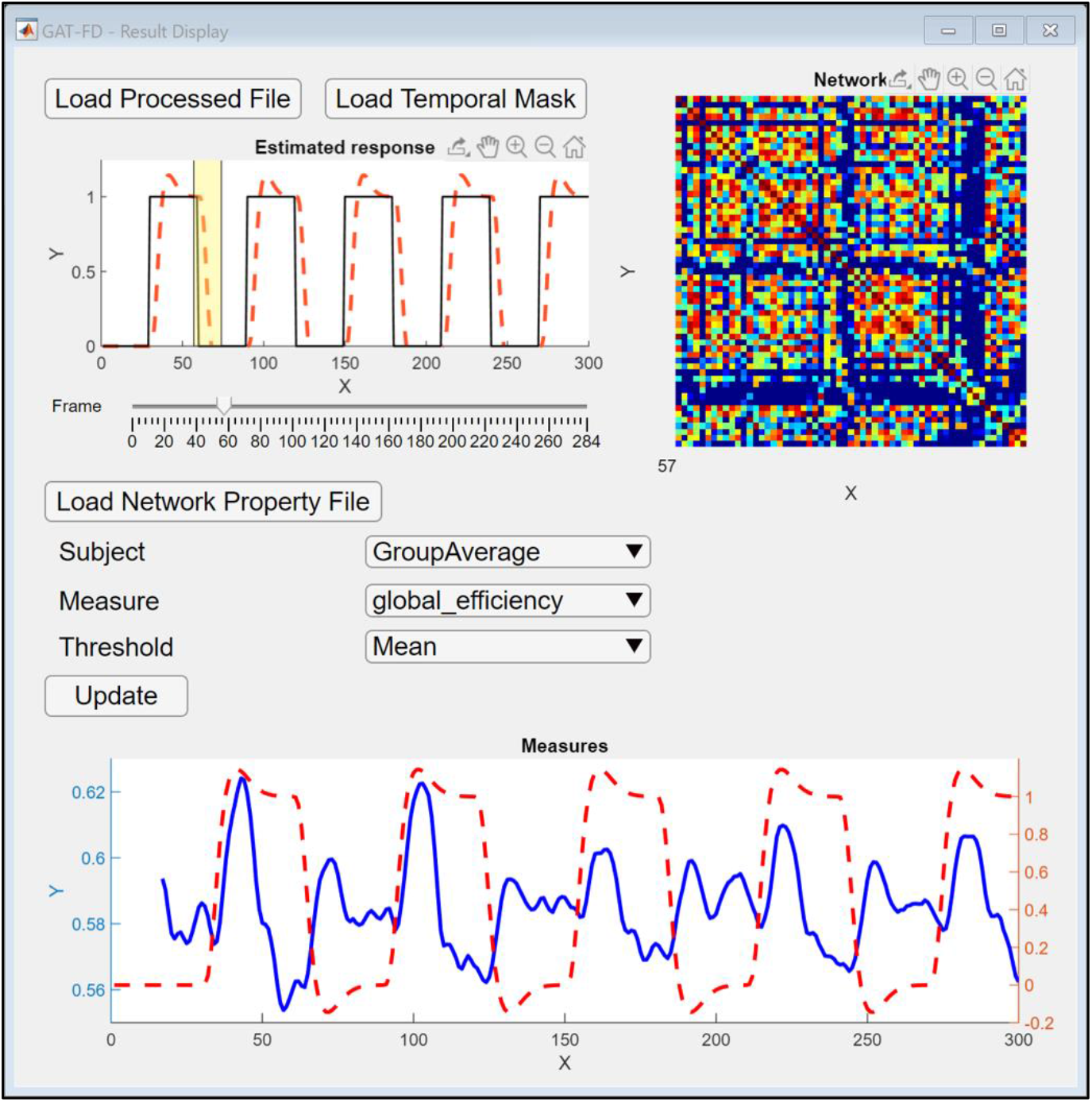
The user interface of result display module. After loading the generated connectivity matrices file and temporal mask file, the functional connectivity matrix can be displayed at top right conner by selecting the desired sliding-window step using the “Frame” slider. After loading the generated network properties file, the network topological properties for different subjects and different threshold values can be displayed at the bottom by clicking the “Update” button.

## 3. Illustration

### 3.1. Data

Task-based fMRI data from 40 typically developing children (male/female: 22/18) were involved in the validation and illustration of the GAT-FD toolbox. All subjects were 11 to 16 years old, right-handed according to the Edinburgh Handedness Inventory (Oldfield, 1971), within or post puberty based on the parent version of Carskadon and Acebo’s rating scale (Carskadon and Acebo, 1993), and had full scale IQ≥ 80 estimated by the Wechsler Abbreviated Scale of Intelligence II (WASI-II) (Wechsler, 2011). None of the subjects reported a history or current diagnoses of neurological and psychiatric disorders, chronic medial illnesses, or learning disabilities. None of them had been taking stimulant or non-stimulant medications within the past 3 months prior to the study visit that might impact the brain activations during fMRI data acquisition. The study received institutional review board approval at the New Jersey Institute of Technology, Rutgers University, and Saint Peter’s University Hospital. Prior the study, all the participants and their parents or legal guardians provided written informed assents and consents, respectively.

Each participant performed a block-design visual sustained attention task (VSAT) during the fMRI scan. The VSAT included 5 task blocks interleaved by 5 rest blocks, each was 30 seconds. Detailed descriptions of the task were provided in our previous studies (Xia et al., 2014;Wu et al., 2018). The fMRI data were collected using a 3-Tesla Siemens TRIO (Siemens Medical Systems, Germany) scanner with a whole brain gradient echo-planar sequence (voxel size = 1.5 mm × 1.5 mm × 2. 0 mm, TR = 1000 ms, echo time = 28.8 ms, and field of view = 208 mm, slice thickness = 2.0 mm).

### 3.2. Preprocessing

The preprocessing steps were performed using FEAT toolbox in FMRIB’s Software Library (Smith et al., 2004). Each raw data was first corrected for slice timing and motion artifacts, using sinc interpolation and rigid-body transformation, respectively. Then brain extraction was performed to remove non-brain tissues using the averaged fMRI data. Spatial smoothing was performed with a 5-mm full-width at half maximum gaussian kernel to improve the signal-to-noise ratio. The signal intensity was then normalized for each slice. Then, a high-pass temporal filter of 1/75 Hz was applied to remove low frequency noises. Finally, linear registration was performed to the MNI template with a voxel size of 2 × 2 × 2 mm^3^.

The group average activation map within the study cohort was calculated and parcellated according to the AAL atlas (Tzourio-Mazoyer et al., 2002). Within each parcellated brain regions that contain at least 100 significantly activated voxels (Z ≥ 2.3 after cluster correction for multiple comparisons), a spherical ROI with the radius of 5mm and centered at the regional maximum of the activated cluster was generated in the MNI space. A total of 59 ROIs (nodes for the to be constructed functional brain networks) were generated and mapped back to each pre-processed fMRI data to construct the dynamic functional networks.

### 3.3. The GAT-FD-based processes

For data from each subject, wavelet-based temporal filtering was first performed on the time series of each ROI, using the 5-level wavelet transformation. The level 3,4, and 5, corresponding to frequency band of 0.015-0.124 Hz, were then used to reconstruct the time series for each ROI. This frequency band has been demonstrated to contain most task-related hemodynamic information (Bassett et al., 2011;Li et al., 2012;Xia et al., 2014). Then, the sliding-window analysis was performed with the window size of 17 TRs and the step size of 1 TR. Such window size and step size were suggested to be able to generate reliable temporal correlation coefficient (the FC measure) within each sliding step and offer enough sliding steps that allows for estimations of variability of the FC during the task period (Di et al., 2015;Li et al., 2018;Betzel et al., 2020). A total of 284 connectivity matrices were then generated for each subject.

As an example of illustrating the characterization the task-related dynamics of the network properties, a temporal inclusive mask was generated based on the blocked design of the task. The condition vector was set as “0 1 0 1 0 1 0 1 0 1”, where 0 was for a rest block and 1 for a task block. The duration vector for each block was set as “30 30 30 30 30 30 30 30 30 30”, with a unit of second. The estimated activation level threshold was set as 0.8, and the task condition coverage percentage threshold was set as 80%, as suggested by default. By implementing this temporal inclusive mask, a total of 79 task-related FC matrices, which were generated in the sliding-window analysis module, were marked as task related.

In the network property analysis module, the absolute thresholding method was implemented for the binarization of the 284 FC matrices generated in the sliding-window analysis module. The range of correlation coefficient threshold was set as from 0.5 to 0.85 with a step size of 0.01. Such threshold range was calculated based on the cost-range of the functional network that satisfied the small-world network assumption (Watts and Strogatz, 1998;Sporns and Honey, 2006). Then the global and nodal topological properties of each of the 284 functional brain networks were calculated. In each subject, the variance of each topological property was calculated over the 284 time points in the overall task duration and over the 79 task-related timepoints based on the generated temporal mask. The group average of network topological properties was then calculated. In addition, variances of the network properties calculated over the full task duration were compared with those calculated based on the generated temporal mask, using paired sample t-test, with a threshold of significance at α≤ 0.05.

## 4. Results and Discussion

As an example of visualization of the generated results, the group mean of the task-related dynamics of the network global efficiency was shown in **Figure 6**. We observed increased global efficiency for information transferring among the 59 nodes in the functional brain network during both rest-to-task and task-to-rest transition periods, while a relatively steady state of this topological property in the middle of the task blocks. Such pattern of task-related FC dynamics has also been observed in other studies (Betti et al., 2013;Cole et al., 2014;Di et al., 2015;Kwon et al., 2017). In addition, all the network topological properties showed significantly lower variances during the task-related period covered by the temporal mask, when compared to those estimated over the entire task duration (the network global efficiency (t=2.282, *p*=0.028), network local efficiency (t=2.223, *p*=0.032), network averaged degree (t=2.529, *p*=0.016), and network averaged clustering coefficient (t=2.235, *p*=0.031)). Indeed, superior topological stability of the functional brain network during task performance relative to that during resting state has also been reported by other studies (Chen et al., 2015;Elton and Gao, 2015).

**Figure 6.**
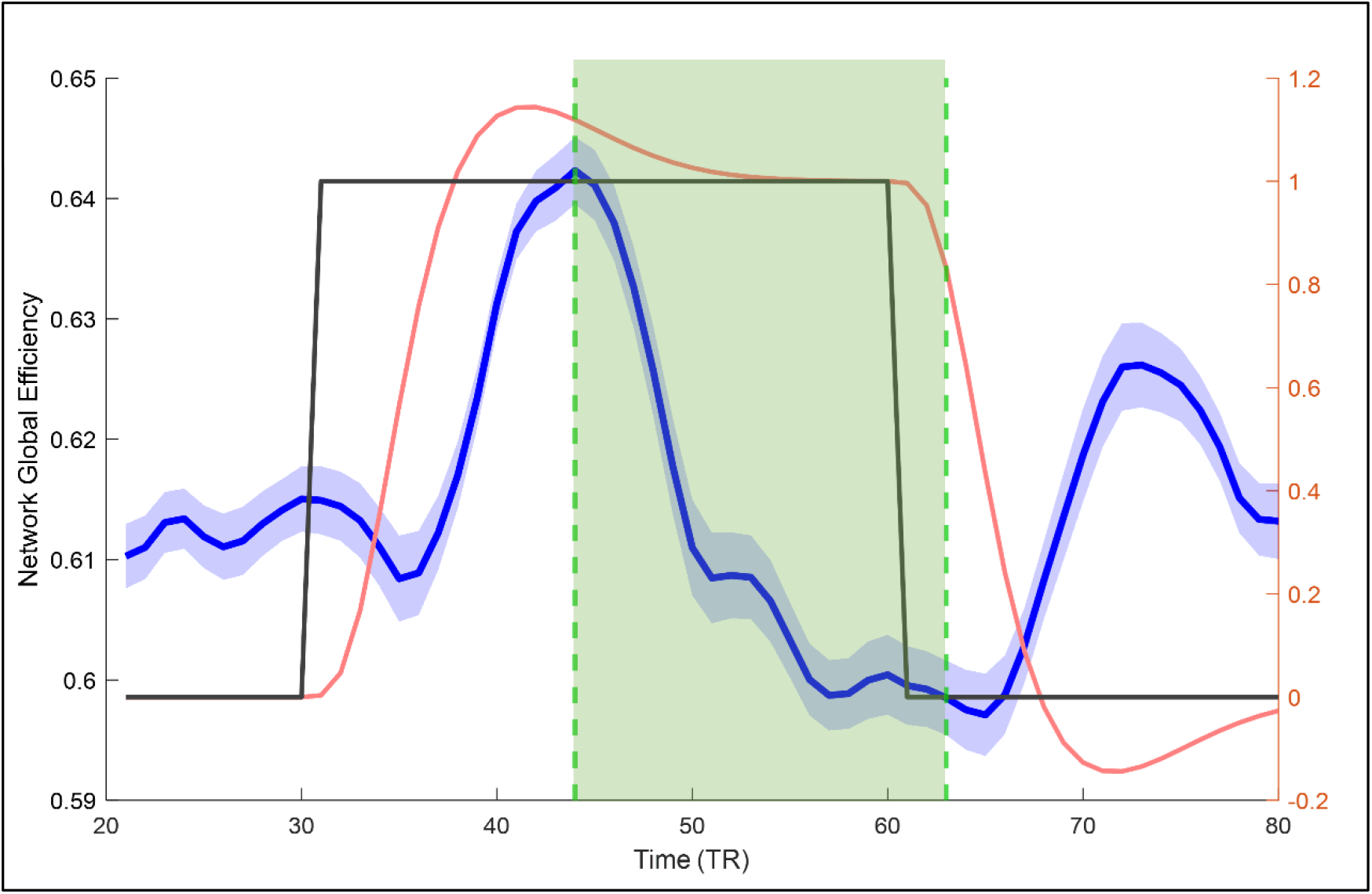
The group average of network global efficiency in response to the experimental task, which were averaged over the first 4 task blocks (the solid blue line). The light blue shadowed area represents the standard error of the mean of the network global efficiency. The task condition is plotted using the solid black line. The estimated hemodynamic response over one testing block is plotted using the solid red line. The task-related temporal inclusive mask is shadowed in light green.

On the bases of the initial pipelines provided in the current version of the GAT-FD toolbox, future work will focus on developing and including more alternative analytical techniques for characterizing the dynamics of FC and network properties.

## 3.2. Conclusion

In this study, we introduced an integrative MATLAB toolbox, GAT-FD, for analyzing the task-related dynamics of FC and topological properties of the functional brain networks for sensory and cognitive processes during task-based fMRI, especially for block-designed data. All the involved functions have been tested and validated using data collected from human subjects during task-based fMRI. The results demonstrated that the GAT-FD allows for effective and quantitative evaluations of the functional network properties and their dynamics during the entire task or user-specified periods. The GAT-FD toolbox and user manual are freely available at https://www.nitrc.org/projects/gat_fd and https://centers.njit.edu/cnnl/gat_fd/.

## 6. Acknowledgements

Specific Acknowledgement goes to the National Institute of Mental Health and the New Jersey Commission on Brain Injury Research for their funding support of this study.

